# Molecular basis of hemoglobin adaptation in the high-flying bar-headed goose

**DOI:** 10.1101/222182

**Authors:** Chandrasekhar Natarajan, Agnieszka Jendroszek, Amit Kumar, Roy E. Weber, Jeremy R. H. Tame, Angela Fago, Jay F. Storz

## Abstract

During adaptive phenotypic evolution, some selectively fixed mutations may be directly causative and others may be purely compensatory. The relative contribution of these two classes of mutation depends on the form and prevalence of mutational pleiotropy. To investigate the nature of adaptive substitutions and their pleiotropic effects, we used a protein engineering approach to characterize the molecular basis of hemoglobin (Hb) adaptation in the bar-headed goose (*Anser indicus*), a hypoxia-tolerant species renowned for its trans-Himalayan migratory flights. We synthesized and tested all possible mutational intermediates in the line of descent connecting the wildtype bar-headed goose genotype with the most recent common ancestor of bar-headed goose and its lowland relatives. Site-directed mutagenesis experiments revealed effect-size distributions of causative mutations and biophysical mechanisms underlying changes in function. Trade-offs between alternative functional properties revealed the importance of compensating deleterious pleiotropic effects in the adaptive evolution of protein function.

## Introduction

During the adaptive evolution of a given trait, some of the selectively fixed mutations will be directly causative (contributing to the adaptive improvement of the trait itself) and some may be purely compensatory (alleviating problems that were created by initial attempts at solution). Little is known about the relative contributions of these two types of substitution in adaptive phenotypic evolution and much depends on the prevalence and magnitude of antagonistic pleiotropy (Burch & Chao, 1999; Cooper, Ostrowski, & Travisano, 2007; Moore, Rozen, & Lenski, 2000; Ostrowski, Rozen, & Lenski, 2005; Otto, 2004; Poon & Chao, 2006; Qian, Ma, Xiao, Wang, & Zhang, 2012; Stern, 2000; Szamecz et al., 2014). If mutations that produce an adaptive improvement in one trait have adverse effects on other traits, then the fixation of such mutations will select for compensatory mutations to mitigate the deleterious side effects, and evolution will proceed as a ‘two steps forward, one step back’ process. In systems where it is possible to identify the complete set of potentially causative mutations that are associated with an adaptive change in phenotype, key insights could be obtained by using reverse genetics experiments to measure the direct effects of individual mutations on the selected phenotype in conjunction with assessments of mutational pleiotropy in the same genetic background.

To investigate the nature of adaptive mutations and their pleiotropic effects, we used a protein engineering approach to characterize the molecular basis of hemoglobin (Hb) adaptation in the high-flying bar-headed goose (*Anser indicus*). This hypoxia-tolerant species is renowned for its trans-Himalayan migratory flights (Hawkes et al., 2011; Hawkes et al., 2013; Bishop et al., 2015), and its elevated Hb-O_2_ affinity is thought to make a key contribution to its capacity for powered flight at extreme elevations of 6000-9000 m (Petschow et al., 1977; Black & Tenney, 1980; Faraci, 1986; Scott & Milsom, 2006; Scott & Milsom, 2007; Scott, 2011; Meir & Milsom, 2013; Scott et al., 2015). At such elevations, an increased Hb-O_2_ affinity helps safeguard arterial O_2_ saturation, thereby compensating for the low O_2_ tension of inspired air. This can help sustain O_2_ delivery to metabolizing tissues because if environmental hypoxia is sufficiently severe, the benefit of increasing pulmonary O_2_ loading typically outweighs the cost associated with a lower O_2_ unloading pressure in the systemic circulation (Bencowitz, Wagner, & West, 1982; Willford, Hill, & Moores, 1982; Storz, 2016).

The Hb of birds and other jawed vertebrates is a heterotetramer consisting of two α-chain and two β-chain subunits. The Hb tetramer undergoes an oxygenation-linked transition in quaternary structure, whereby the two semi-rigid α_1_β_1_ and α_2_β_2_ dimers rotate around one another by 15° during the reversible switch between the deoxy (low-affinity [T]) conformation and the oxy (high-affinity [R]) conformation (Perutz, 1972; Baldwin & Chothia, 1979; Lukin & Ho, 2004; Yuan, Tam, Simplaceanu, & Ho, 2015). Oxygenation-linked shifts in the T↔R equilibrium govern the cooperativity of O_2_-binding and are central to Hb’s role in respiratory gas transport.

The major Hb isoform of the bar-headed goose has an appreciably higher O_2_-affinity than that of the closely related greylag goose (*Anser anser*), a strictly lowland species (Petschow et al., 1977; Rollema & Bauer, 1979). The Hbs of the two species differ at five amino acid sites: three in the α*^A^*-chain subunit and two in the β*^A^*-chain subunit (Oberthur, Braunitzer, & Wurdinger, 1982; McCracken, Barger, & Sorenson, 2010). Of these five amino acid differences, Perutz (Perutz, 1983) predicted that the Pro→Ala replacement at α119 (αP119A) is primarily responsible for the adaptive increase in Hb-O_2_ affinity in bar-headed goose. This site is located at an intersubunit (α_1_β_1_/α_2_β_2_) interface where the ancestral α119-Pro forms a van der Waals contact with β55-Met on the opposing subunit of the same αβ dimer. Perutz predicted that the αP119A mutation would eliminate this intradimer contact, thereby destabilizing the T-state and shifting the conformational equilibrium in favor of the high-affinity R-state. Jessen et al. (Jessen, Weber, Fermi, Tame, & Braunitzer, 1991) and Weber et al. (Weber, Jessen, Malte, & Tame, 1993) tested Perutz’s hypothesis using a protein engineering approach based on site-directed mutagenesis, and their experiments confirmed the predicted mechanism.

As a result of these experiments, bar-headed goose Hb is often held up as an example of a biochemical adaptation that is attributable to a single, large-effect substitution (Li, 1997; Hochachka & Somero, 2002). However, several key questions remain unanswered: Do the other substitutions also contribute to the change in Hb-O_2_ affinity? If not, do they compensate for deleterious pleiotropic effects of the affinity-enhancing αP119A mutation? Given that the substitutions in question involve closely linked sites in the same gene, another possibility is that neutral mutations at the other sites simply hitchhiked to fixation along with the positively selected mutation. Since the other mutations in bar-headed goose Hb have not been tested, we do not know whether αP119A accounts for all or most of the evolved change in O_2_ affinity. Moreover, the original studies tested the effect of αP119A by introducing the goose-specific amino acid state into recombinant human Hb. One potential problem with this type of ‘horizontal’ comparison – where residues are swapped between orthologous proteins of contemporary species – is that the focal mutation is introduced into a sequence context that is not evolutionarily relevant. If mutations have context-dependent effects, then introducing goose-specific substitutions into human Hb may not recapitulate the phenotypic effects of the mutations on the genetic background in which they actually occurred (i.e., in the ancestor of bar-headed goose). An alternative ‘vertical’ approach is to reconstruct and resurrect ancestral proteins to test the effects of historical mutations on the genetic background in which they actually occurred during evolution (Harms & Thornton, 2010; Hochberg & Thornton, 2017).

Here we revisit the functional evolution of bar-headed goose Hb, a classic text-book example of biochemical adaptation. We reconstructed the α*^A^* - and β*^A^*-chain Hb sequences of the most recent common ancestor of the bar-headed goose and its closest living relatives, all of which are lowland species in the genus *Anser*. After identifying the particular substitutions that are specific to bar-headed goose, we used a combinatorial approach to test the functional effects of each mutation in all possible multi-site combinations. To examine possible pleiotropic effects of causative mutations, we also measured several properties that potentially trade-off with Hb-O_2_ affinity: susceptibility to spontaneous heme oxidation (autoxidation rate), allosteric regulatory capacity (the sensitivity of Hb-O_2_ affinity to modulation by anionic effectors), and measures of both secondary and tertiary structural stability. Measuring the direct and indirect effects of these mutations enabled us to address two fundamental questions about molecular adaptation: (*i*) Do each of the mutations contribute to the increased Hb-O_2_ affinity? If so, what are their relative effects? And (*ii*) Do function-altering mutations have deleterious pleiotropic effects on other aspects of protein structure or function? If so, are these effects compensated by mutations at other sites?

## Results and Discussion

### Direction of amino acid substitutions

Using globin sequences from bar-headed goose, greylag goose, and other waterfowl species in the subfamily Anserinae, we reconstructed the α- and β-chain sequences of the bar-headed goose/greylag goose ancestor, which we call ‘AncAnser’ because it represents the most recent common ancestor of all extant species in the genus *Anser* (*Figure 1A*). The principle of parsimony clearly indicates that all three of the α-chain substitutions that distinguish the Hbs of bar-headed goose and greylag goose occurred in the bar-headed goose lineage (Gα18S, Aα63V, and αP119A), whereas each of the two β-globin substitutions occurred in the greylag goose lineage (βT4S and βD125E)(*Figures 1A,B*).

**Figure 1.**
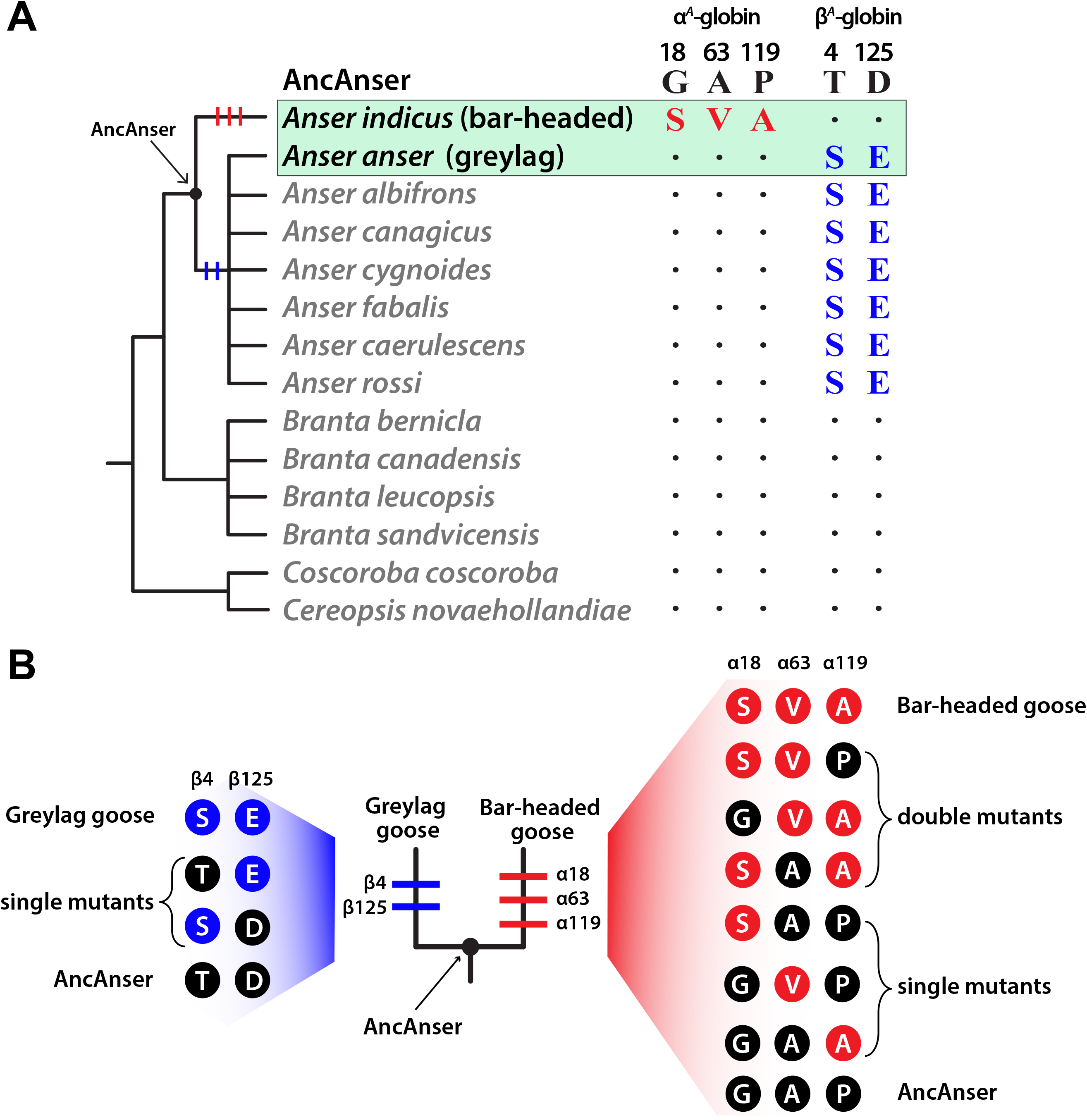
Inferred history of amino acid substitution at five sites that distinguish the major Hb isoforms of the bar-headed goose (*Anser indicus*) and greylag goose (*Anser anser*). (*A*) Amino acid states at the same sites are shown for 12 other waterfowl species in the subfamily Anserinae. Of the five amino acid substitutions that distinguish the Hbs of *A. indicus* and *A. anser*, parsimony indicates that three occurred on the branch leading to *A. indicus* (αG18S, αA63V, and αP119A) and two occurred on the branch subtending the clade of all *Anser* species other than *A. indicus* (βT4S and βD125E). ‘AncAnser’ represents the reconstructed sequence of the *A. indicus/A. anser* common ancestor, which is also the most recent common ancestor of all extant species in the genus *Anser*. (*B*) Triangulated comparisons involving rHbs of bar-headed goose, greylag goose, and their reconstructed ancestor (AncAnser) reveal the polarity of changes in character state. Differences in Hb function between bar-headed goose and AncAnser reflect the net effect of three substitutions (αG18S, αA63V, and αP119A) and differences between greylag goose and AncAnser reflect the net effect of two substitutions (βT4S and βD125E). All possible mutational intermediates connecting AncAnser with each of the two descendent species are shown to the side of each terminal branch.

### Ancestral protein resurrection and functional testing

It is often implicitly assumed that the difference in Hb-O_2_ affinity between bar-headed goose and greylag goose is attributable to a derived increase in Hb-O_2_ affinity in the bar-headed goose lineage (Black & Tenney, 1980; Gillespie, 1991; Li, 1997; Hochachka & Somero, 2002). In principle, however, the pattern could be at least partly attributable to a derived reduction in Hb-O_2_ affinity in the greylag goose lineage, even if αP119A does account for the majority of the change in bar-headed goose. To resolve the polarity of character state change, we synthesized, purified, and functionally tested recombinant Hbs (rHbs) representing the wildtype Hb of bar-headed goose, the wildtype Hb of greylag goose, and the reconstructed Hb of their common ancestor, AncAnser. Functional differences between bar-headed goose and AncAnser rHbs reflect the net effect of three substitutions (αG18S, αA63V, and αP119A) and differences between greylag goose and AncAnser reflect the net effect of two substitutions (βT4S and βD125E; *Figure 1B*).

Since genetically based differences in Hb-O_2_ affinity may be attributable to differences in intrinsic O_2_-affinity and/or changes in sensitivity to allosteric effectors in the red blood cell, we measured O_2_-equilibria of purified rHbs under four standardized treatments: (*i*) in the absence of allosteric effectors (stripped), (*ii*) in the presence of Cl^−^ ions (added as KCl), (*iii*) in the presence of inositol hexaphosphate (IHP, a chemical analog of the endogenously produced inositol pentaphosphate), and (*iv*) in the simultaneous presence of KCl and IHP. This latter treatment is most relevant to *in vivo* conditions in avian red blood cells. In each treatment, we measured *P*_50_ (the partial pressure of O_2_ [PO_2_] at which Hb is 50% saturated). To complement equilibrium measurements on the set of three rHbs and to gain further insight into functional mechanisms, we also performed stopped-flow kinetic experiments to estimate O_2_ binding and dissociation rates under the same conditions.

The O_2_-equilibrium measurements confirmed the results of previous studies (Petschow et al., 1977; Rollema& Bauer, 1979) by demonstrating that the Hb of bar-headed goose has a higher intrinsic O_2_-affinity than that of greylag goose (as revealed by the lower *P*_50_ for stripped Hb)(Figure 2A, *Table 1*). This difference persisted in the presence of Cl^−^ ions (*P*_50(KCl)_), in the presence of IHP (*P*_50(IHP)_), and in the simultaneous presence of both anions (*P*_50(KCl+IHP)_) (*Figure 2A, Table 1*). The difference in Hb-O_2_ affinity between bar-headed goose and greylag goose is mainly attributable to differences in intrinsic affinity, as there were no appreciable differences in sensitivities to allosteric effectors (Table 1). This is consistent with a previous report that native Hbs of bar-headed goose and greylag goose have identical binding constants for inositol pentaphosphate (Rollema& Bauer, 1979). Pairwise comparisons between each of the two modern-day species and their reconstructed ancestor (AncAnser) revealed that the elevated Hb-O_2_ affinity of the bar-headed goose is a derived character state. O_2_-equilibrium properties of greylag goose and AncAnser rHbs were generally very similar (*Figure 2A*). The triangulated comparison involving rHbs from the two contemporary species (bar-headed goose and greylag goose) and their reconstructed ancestor (AncAnser) revealed that – in the presence of physiological concentrations of Cl^−^ and IHP – 72% of the difference in Hb-O_2_ affinity between bar-headed goose and greylag goose is attributable to a derived increase in the bar-headed goose lineage and the remaining 28% is attributable to a derived reduction in the greylag goose lineage (*Figure 2A*). This demonstrates the value of ancestral protein resurrection for inferring the direction and magnitude of historical changes in character state.

**Figure 2.**
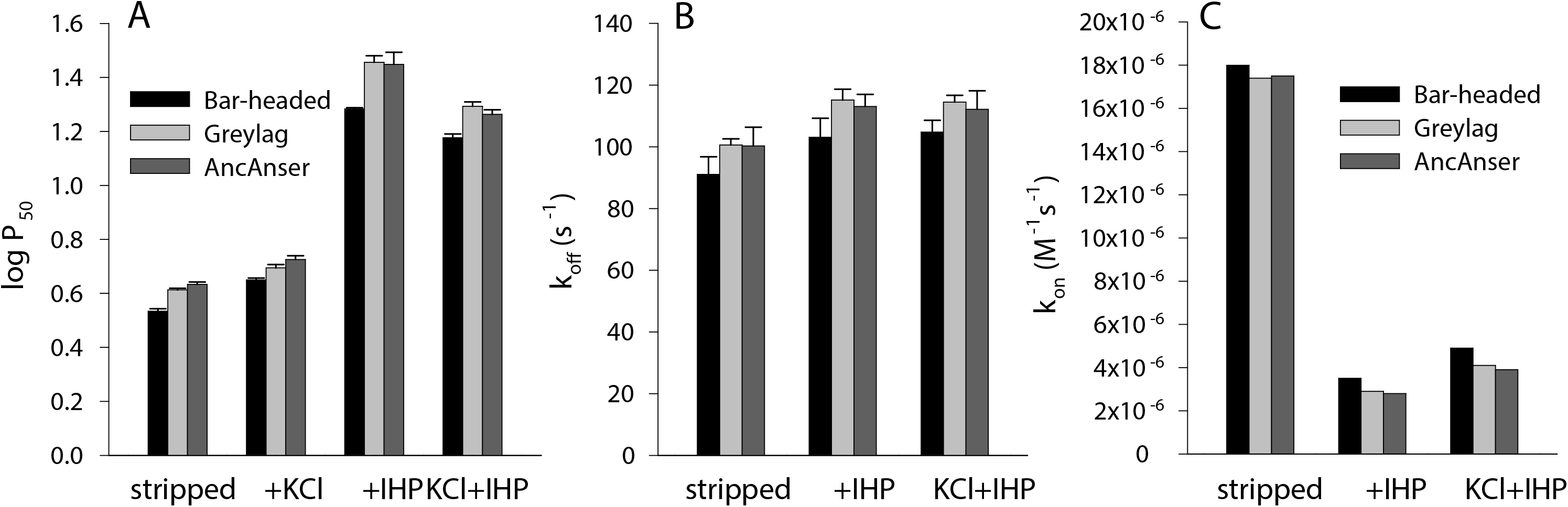
Bar-headed goose evolved a significant increase in Hb-O_2_ affinity relative to greylag goose and their reconstructed ancestor, AncAnser. Triangulated comparisons of purified rHbs involved diffusion-chamber measurements of O_2_-equilibria (*A*) and stopped-flow measurements of O_2_ dissociation kinetics (*B*). O_2_-association rate constants (*k*_on_, M^−1^s^−1^) derived from data in (*A*) and (*B*) are shown in (*C*). O_2_-affinities (*P*_50_, torr; ± 1 SE) and dissociation rates (*k*_off_, M^−1^s^−1^) of purified rHbs were measured at pH 7.4, 37° C, in the absence (stripped) and presence of allosteric effectors ([Cl^−^], 0.1 M; [Hepes], 0.1 M; IHP/Hb tetramer ratio = 2.0; [heme], 0.3 mM in equilibrium experiments; [Cl^−^], 1.65 mM; [Hepes], 200 mM; IHP/Hb tetramer ratio =2.0; [heme], 5 μM in kinetic experiments).

**Table 1.**
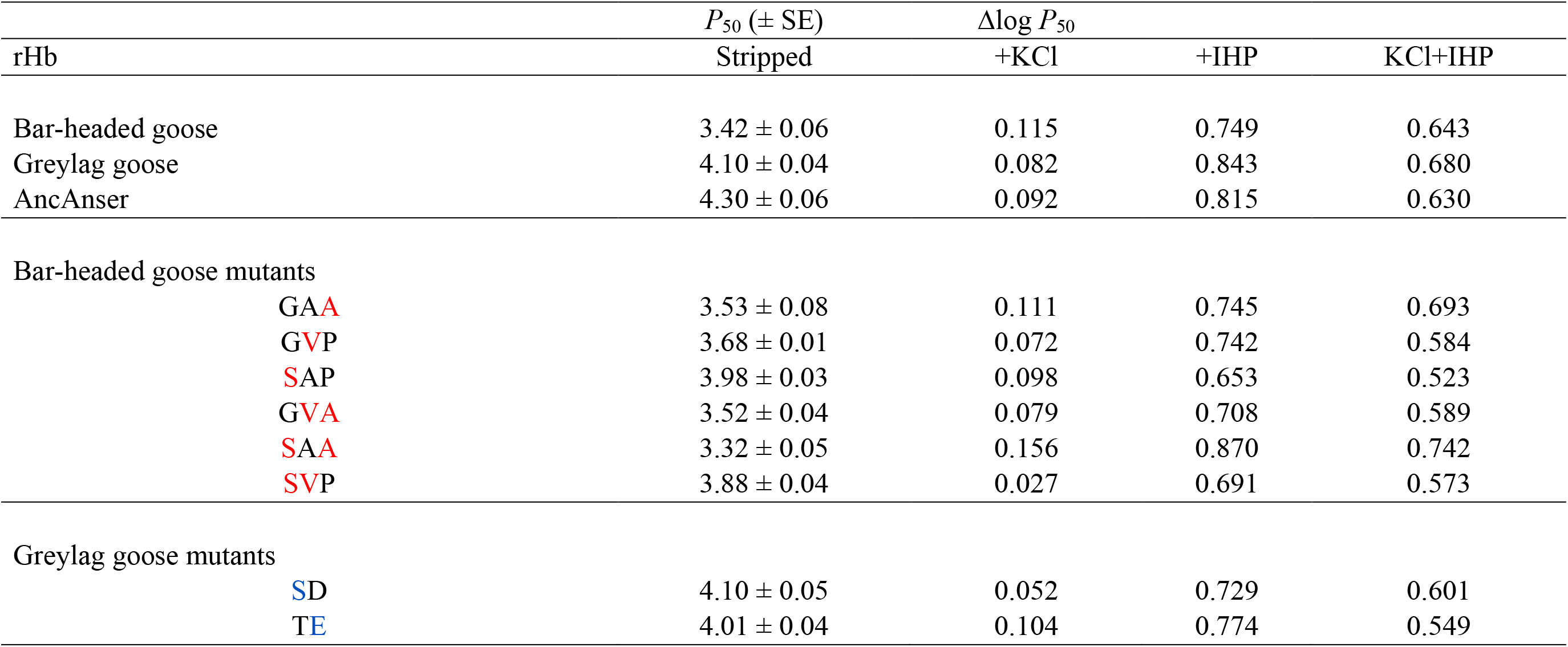
O_2_ affinities (*P*_50_, torr) and anion sensitivities (Δlog *P*_50_) of rHbs representing bar-headed goose, greylag goose, their reconstructed ancestor (AncAnser), and all possible mutational intermediates connecting AncAnser with each of the two descendant species. O_2_ equilibria were measured in 0.1 mM Hepes buffer at pH 7.4 (± 0.01) and 37°C in the absence (stripped) and presence of Cl^−^ ions (0.1 M KCl]) and IHP (at two-fold molar excess over tetrameric Hb). Anion sensitivities are indexed by the difference in log-transformed values of *P*_50_ in the presence and absence of Cl^−^ ions (KCl) and IHP. The higher the Δlog *P*_50_ value, the higher the sensitivity of Hb-O_2_ affinity to the presence of a given anion or combination of anions. For the bar-headed goose mutants (all mutational intermediates between wildtype bar-headed goose and AncAnser), three-letter genotype codes denote amino acid states at α18, α63, and α119 (amino acid abbreviations in black lettering = ancestral, red lettering = derived). At these same three sites, AncAnser is ‘GAP’ the wildtype genotype of bar-headed goose is ‘SVA’. For the greylag goose mutants (all mutational intermediates between wildtype greylag goose and AncAnser), two-letter genotype codes denote amino acid states at β4 and β125 (amino acid abbreviations in black lettering = ancestral, blue lettering= derived). At these same two sites, AncAnser is ‘TD’ the wildtype genotype of greylag goose is ‘SE’.

Kinetic measurements demonstrated that the increased O_2_-affinity of bar-headed goose rHb (i.e., the lower ratio of dissociation/association rate constants, *k*_off_/*k*_on_) is attributable to a lower rate of dissociation, *k*_off_, in combination with a faster rate of O_2_-binding, *k*_on_, relative to the Hbs of both greylag goose and AncAnser (*Figure 2B,C*). The Hbs of greylag goose and AncAnser exhibited highly similar rates of both *k*_off_ and *k*_on_(*Figure 2B,C*).

### Effects of individual mutations in bar-headed goose Hb

In combination with the inferred history of sequence changes (*Figure 1A,B*), the comparison between the rHbs of bar-headed goose and AncAnser indicates that the derived increase in Hb-O_2_ affinity in barheaded goose must be attributable to the independent or joint effects of the three substitutions at sites α18, α63, and α119. To measure the effects of each individual mutation in all possible multi-site combinations, we used site-directed mutagenesis to synthesize each of the six possible mutational intermediates that connect the ancestral and descendant genotypes (*Figure 1B*). In similar fashion, we synthesized each of the two possible mutational intermediates that connect AncAnser and the wildtype genotype of greylag goose (*Figure 1B*).

The analysis of the bar-headed goose mutations on the AncAnser background revealed that mutations at each of the three α-chain sites (αG18S, αA63V, and αP119A) produced significant increases in intrinsic Hb-O_2_ affinity (indicated by reductions in *P*_50(stripped)_)(*Figure 3, Table 1*). The Pα119A mutation had the largest effect on the ancestral background, producing an 18% reduction in *P*_50(stripped)_ (increase in intrinsic Hb-O_2_ affinity). On the same background, αG18S or αA63V produced 7% and 14% reductions in *P*_50(stripped)_, respectively. In the set of six (=3!) possible mutational pathways connecting the low-affinity AncAnser genotype (GAP) and the high-affinity bar-headed goose genotype (SVA), the αP119A mutation produced a significant increase in Hb-O_2_ affinity on each of four possible backgrounds (corresponding to the first step in the pathway, two alternative second steps, and the third step; *Figure 3*). When tested on identical backgrounds, αP119A invariably produced a larger increase in intrinsic Hb-O_2_ affinity than either αG18S or αA63V. Nonetheless, of the six possible forward pathways connecting GAP and SVA, αP119A had the largest effect in four pathways and αA63V had the largest effect in the remaining two. The two pathways in which αA63V had the largest effect were those in which it occurred as the first step. In fact, αG18S or αA63V only produced significant increases in Hb-O_2_ affinity when they occurred as the first step. The effects of these two mutations were always smaller in magnitude when they occurred on backgrounds in which the derived α119-Ala was present. In addition to differences in average effect size, αP119A also exhibited a higher degree of additivity across backgrounds than the other two mutations. For example, the affinity-enhancing effect ofαP119A on the AncAnser background is mirrored by a similarly pronounced reduction in O_2_-affinity when the mutation is reverted on the wildtype bar-headed goose background (αA119P). By contrast, forward and reverse mutations at α18 and α63 do not show the same symmetry of effect (*Figure 3 – figure supplement 1*).

**Figure 3.**
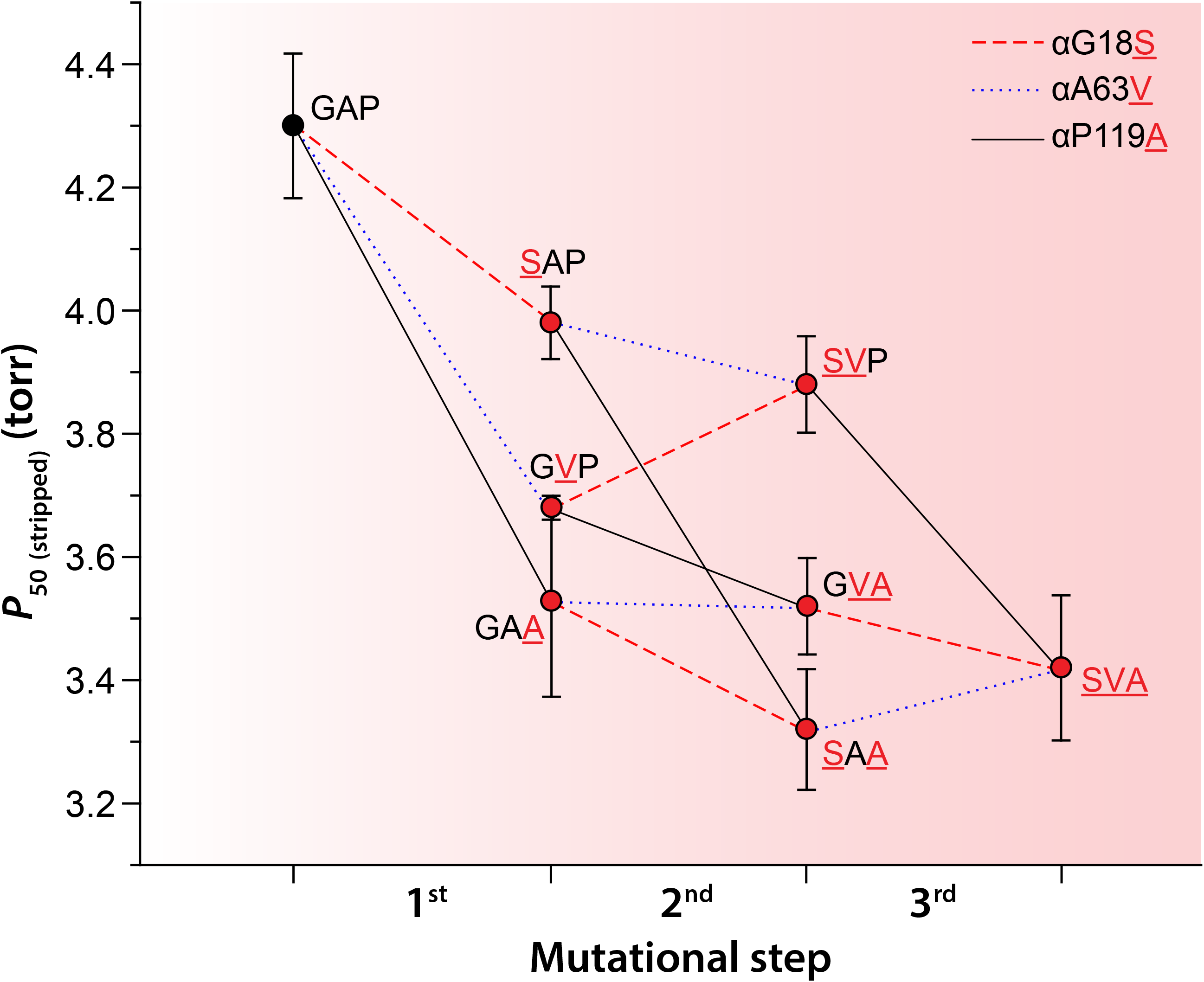
Trajectories of change in intrinsic Hb-O_2_ affinity (indexed by *P*_50_, torr) in each of six possible forward pathways that connect the ancestral ‘AncAnser’ genotype (GAP) and the wildtype genotype of bar-headed goose (SVA). Derived amino acid states are indicated by red lettering. Error bars denote 95% confidence intervals.

### Structural mechanisms underlying the evolved increase in Hb-O_2_ affinity in bar-headed goose

Comparison of crystal structures for human and bar-headed goose Hbs (Liang, Hua, Liang, Xu, & Lu, 2001) revealed that each of the three bar-headed goose α-chain substitutions have structurally localized effects. As noted by Jessen et al. (Jessen et al., 1991), the αP119A mutation has very little effect on the main-chain formation, and appears to exert its functional effect via the elimination of side chain contacts and increased backbone flexibility. The αA63V mutation is predicted to increase flexibility in the ‘AB corner’ (formed by the juncture between the A and B helices), as the introduction of the valine side chain causes minor steric clashes with two neighboring glycines at α-chain sites 25 and 59 (*Figure 4*). This interaction may increase O_2_-affinity by impinging on the neighboring α58-His, the ‘distal histidine’ that stabilizes the α-heme Fe-O_2_ bond (Olson et al., 1988; Mathews et al., 1989; Rohlfs et al., 1990; Lukin, Simplaceanu, Zou, Ho, & Ho, 2000; Birukou, Schweers, & Olson, 2010; Yuan, Simplaceanu, Ho, & Ho, 2010).

**Figure 4.**
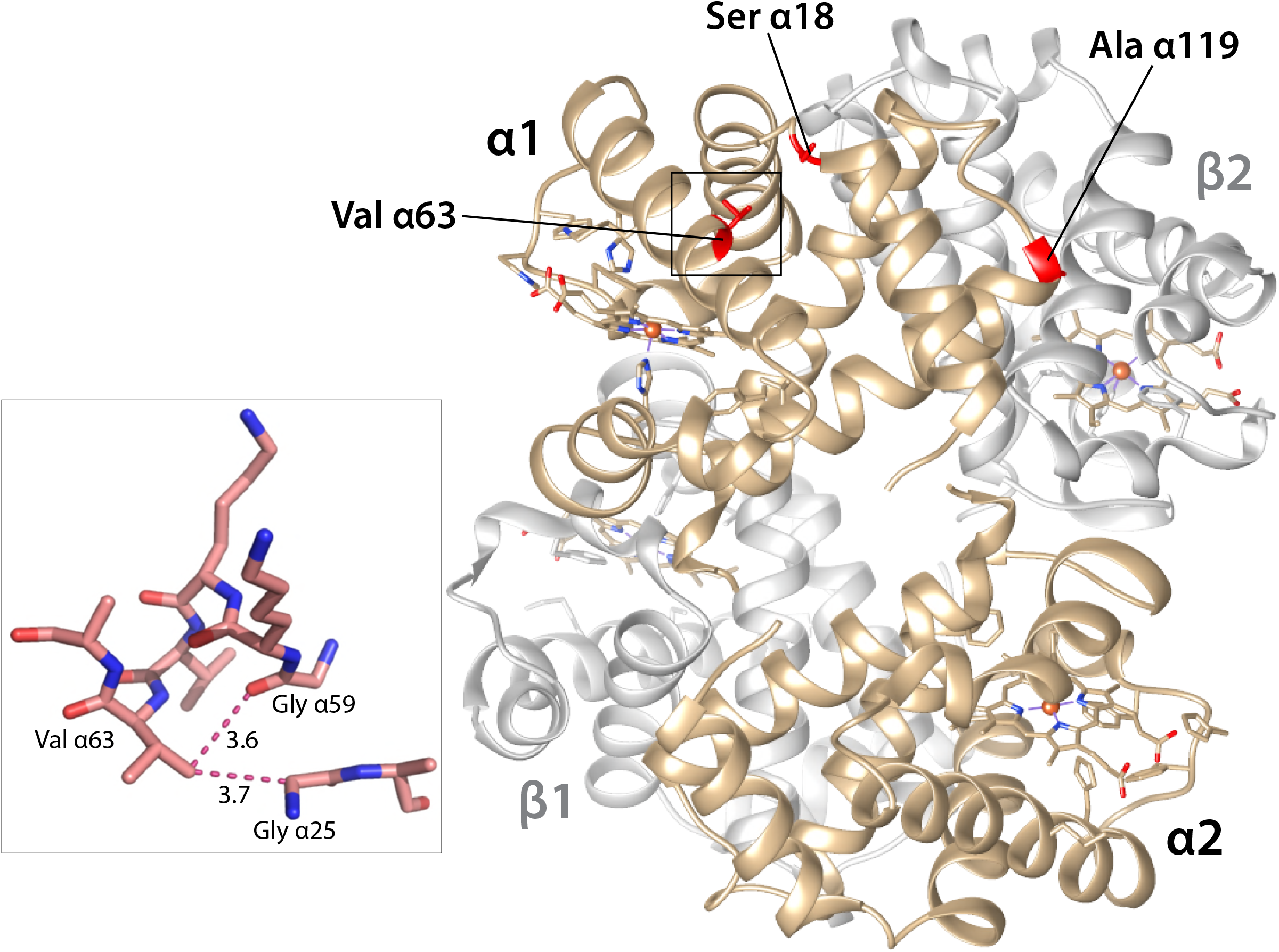
Structural model showing bar-headed goose Hb in the deoxy state (PDB1hv4), along with locations of each of the three amino substitutions that occurred in the bar-headed goose lineage after divergence from the common ancestor of other *Anser* species. The inset graphic shows the environment of the α63-Val residue. When valine replaces the ancestral alanine at this position, the larger volume of the side-chain causes minor steric clashes with two neighboring glycine residues, α25-Gly and α59-Gly and is predicted to increase flexibility of the AB corner. The distances between non-hydrogen atoms (depicted by dotted lines) are given in Å.

### Effects of individual mutations in greylag goose Hb

Given that the AncAnser and greylag goose rHbs exhibit similar equilibrium and kinetic O_2_-binding properties (*Figure 2A,B,C*), the two greylag goose mutations (βT4S and βD125E) obviously do not produce an appreciable net change in combination. Interestingly, however, each mutation by itself produces a slightly reduced sensitivity to IHP (*Table 1*), such that values of *P*_50(IHP)_ and *P*_50(kC1+IHP)_ for the single-mutant intermediates were significantly lower than those for AncAnser and the wildtype genotype of greylag goose.

### Mutational pleiotropy

Since amino acid mutations often affect multiple aspects of protein biochemistry (DePristo, Weinreich, & Hartl, 2005; Harms & Thornton, 2013; Tokuriki et al., 2012; Tokuriki, Stricher, Serrano, & Tawfik, 2008), it is of interest to test whether adaptive mutations that improve one aspect of protein function simultaneously compromise other functions. Amino acid mutations that alter the oxygenation properties of Hb often have pleiotropic effects on allosteric regulatory capacity, structural stability, and susceptibility to heme loss and/or heme oxidation (Kim et al., 1994; Olson, Eich, Smith, Warren, & Knowles, 1997; Olson& Maillett, 2005; Bellelli, Brunori, Miele, Panetta, & Vallone, 2006; Bonaventura, Henkens, Alayash, Banerjee, & Crumbliss, 2013; Tam et al., 2013; Varnado et al., 2013; Kumar et al., 2017). Accordingly, we tested whether mutational changes in intrinsic O_2_-affinity are associated with potentially deleterious changes in other structural and functional properties.

Analysis of the full set of bar-headed goose and greylag goose rHb mutants revealed modest variability in autoxidation rate (*Figure 3 – figure supplement 2A*). This property is physiologically relevant because oxidation of the ferrous (Fe^2+^) heme iron to the ferric state (Fe^3+^) releases superoxide (O_2_^−^) or perhydroxy (HO_2_•) radical, and prevents reversible Fe-O_2_ binding, rendering Hb inoperative as an O_2_-transport molecule. Although mutational changes in intrinsic O_2_ affinity (Δlog *P*_50(stripped)_) were not significantly correlated with changes in autoxidation rate in the full dataset (*r* = −0.311), analysis of the bar-headed goose rHb mutants revealed a striking pairwise interaction between mutations at α18 and α63 (residues which are located within 7 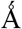 of one another). The αA63V mutation produced a >2-fold increase in the autoxidation rate on backgrounds in which the ancestral Gly is present at α18 (*Figure 5, Figure 3 – figure supplement 2A, source data 1*). The adjacent α62-Val is highly conserved because it plays a critical role in restricting solvent access to the distal heme pocket, thereby preventing water-catalyzed rupture of the Fe-O_2_ bond to release a superoxide ion (Egeberg et al., 1990; Tame, Shih, Pagnier, Fermi, & Nagai, 1991; Quillin et al., 1995; Tam et al., 2013). An increase in side chain volume at α63 may compromise this gating function, resulting in an increased susceptibility to heme oxidation. The increased autoxidation rate caused by αA63V is fully compensated by αG18S (*Figure* 5), a highly unusual amino acid replacement because glycine is the only amino acid at this site (the C-terminal end of the A helix) that permits the main chain to adopt the typical Ramachandran angles (*Figure 4 – figure supplement 1*). Introduction of the serine side chain at α18 in bar-headed goose Hb forces this residue to undergo a peptide flip relative to human Hb, so the carbonyl oxygen points in the opposite direction. This unusual replacement at α18 may be required to accommodate the bulkier Val side chain at α63, thereby alleviating conformational stress.

**Figure 5.**
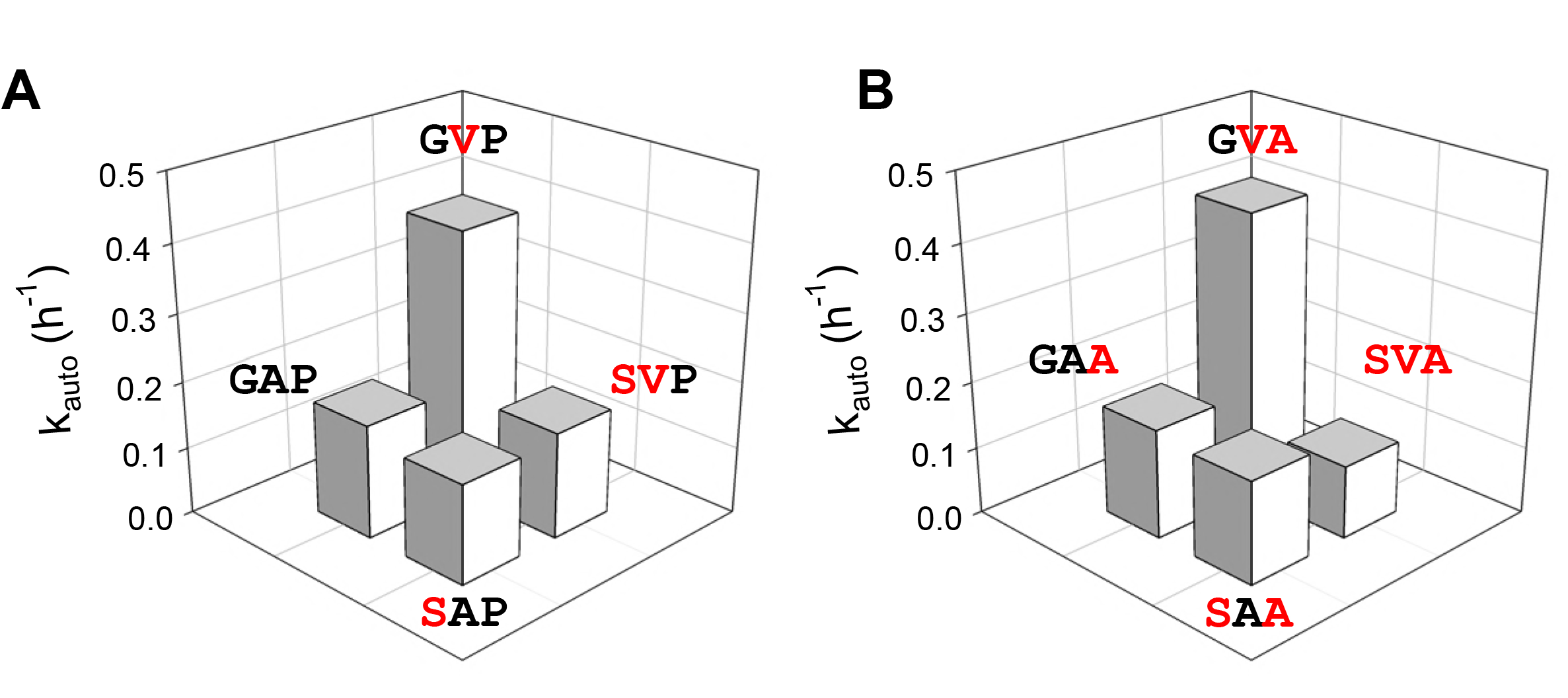
Compensatory interaction between spatially proximal α-chain residues in bar-headed goose Hb. The mutation αA63V produces a >2-fold increase in autoxidation rate (*k*_auto_) on genetic backgrounds with the ancestral Gly at residue position α18. This effect is fully compensated by αG18S, as indicated by two double-mutant cycles (*A* and *B*) in which mutations at both sites are tested individually and in pairwise combination. Derived amino acid states are indicated by red lettering.

Aside from the compensatory interaction between mutations at α18 and α63, we observed no evidence for trade-offs between O_2_-affinity and any of the other measured functional or structural properties. There were no significant correlations between Δlog *P*_50(stripped)_ and changes in allosteric regulatory capacity (*Table* 1), as measured by sensitivity to Cl^−^ (*r* = −0.534), IHP (*r* = −0.137), or both anions in combination (*r* = −0.300). The goose rHbs revealed no appreciable variation in the stability of α- helical secondary structure as measured by circular dichroism spectroscopy (*Figure 3 – figure supplement 2B, source data 2*) and there were no significant correlations between Δlog *P*_50(stripped)_ and changes in stability over the physiological range (pH 6.5, *r* = −0.357; pH 7.5, *r* = −0.052). Likewise, the rHbs exhibited very little variation in the stability of tertiary structure as measured by UV-visible spectroscopy (*Figure 3 – figure supplement 2C, source data 3*) and there were no significant correlations between Δlog *P*_50(stripped)_ and changes in stability over the physiological range (pH 6.5, *r* = −0.511; pH 7.5, *r* = −0.338). In summary, we found no evidence for pleiotropic trade-offs between intrinsic O_2_-affinity and any measured properties of Hb structure or function other than autoxidation rate.

## Conclusions

We now return to the two questions we posed at the outset:

(1) *Do each of the bar-headed goose mutations contribute to the increased Hb-O_2_ affinity?* It depends on the order in which the substitutions occur. Our experiments demonstrated that the αP119A mutation always produced a significant increase in intrinsic Hb-O_2_ affinity regardless of the background in which it occurred. As documented previously (Jessen et al., 1991; Weber et al., 1993), the αP119A mutation also produces a significant affinity-enhancing effect on the far more divergent background of human Hb (which differs from bar-headed goose Hb at 89 of 267 amino acid sites in each αβ half-molecule [33% divergence in protein sequence]). By contrast, αG18S or αA63V only produced significant affinity-enhancing effects when they occurred as the first step in the pathway (on the AncAnser background). If it was advantageous for the ancestor of today’s bar-headed geese to have an increased Hb-O_2_ affinity, our experiments suggest that any of the three α-chain mutations alone would have conferred a beneficial effect, but only αP119A would have produced the same effect after the other two had already fixed. This illustrates an important point about distributions of mutational effect sizes in adaptive walks: in the presence of epistasis, relative effect sizes may be highly dependent on the sequential order in which the substitutions occur.
(2) *Do function-altering mutations have deleterious pleiotropic effects on other aspects of protein structure or function?* On the AncAnser background, the affinity-enhancing mutation, αA63V, produces a pronounced increase in the autoxidation rate. This is consistent with the fact that engineered Hb and myoglobin mutants with altered affinities often exhibit increased autoxidation rates (Brantley, Smerdon, Wilkinson, Singleton, & Olson, 1993; Olson et al., 1997; Tam et al., 2013; Varnado et al., 2013). In the case of bar-headed goose Hb, the increased autoxidation rate caused by αA63V is completely compensated by a polarity-changing mutation at a spatially proximal site, αG18S. This compensatory interaction suggests that the αG18S mutation may have been fixed by selection not because it produced a beneficial main effect on Hb-O_2_ affinity, but because it mitigated the deleterious pleiotropic effects of the affinity-altering αA63V mutation. Alternatively, if αG18S preceded αA63V during the evolution of bar-headed goose Hb, then the (conditionally) deleterious side effects of αA63V would not have been manifest.

Our experiments revealed no evidence to suggest that the affinity-altering αP1119A mutation perturbed other structural and functional properties of Hb. Data on natural and engineered human Hb mutants have provided important insights into structure-function relationships and the nature of trade-offs between different functional properties (Kim et al., 1994; Olson et al., 1997; Bellelli et al., 2006; Steinberg& Nagel, 2009; Tam et al., 2013; Varnado et al., 2013). An important question concerns the extent to which function-altering spontaneous mutations are generally representative of those that eventually fix and contribute to divergence in protein function between species. There are good reasons to expect that the spectrum of pleiotropic effects among spontaneous mutations or low-frequency variants may be different from the spectrum of effects among evolutionary substitutions (mutations that passed through the filter of purifying selection and eventually increased to a frequency of 1.0)(Streisfeld& Rausher, 2011). The affinity-altering mutations that are most likely to fix (whether due to drift or positive selection) may be those that have minimal pleiotropic effects and therefore do not require compensatory mutations at other sites.

## Materials and methods

### Sequence data

We took sequence data for the α*^A^*- and β*^A^*-globin genes of all waterfowl species from published sources (Oberthur et al., 1982; McCracken et al., 2010).

### Vector construction and site-directed mutagenesis

After optimizing nucleotide sequences of AncAnser α*^A^*- and β*^A^*-globin genes in accordance with *E. coli* codon preferences, we synthesized the α*^A^*-β*^A^*-globin cassette (Eurofins MWG Operon). We cloned the globin cassette into a custom pGM vector system (Shen et al., 1993; Natarajan et al., 2011), as described previously (Natarajan et al., 2013; Projecto-Garcia et al., 2013; Cheviron et al., 2014; Galen et al., 2015; Natarajan et al., 2015; Tufts et al., 2015; Natarajan et al., 2016), and we then used site-directed mutagenesis to derive globin sequences of greylag goose, bar-headed goose, and each of the mutational intermediates connecting these wildtype sequences with AncAnser. We conducted the codon mutagenesis using the QuikChange II XL Site-Directed Mutagenesis kit (Agilent Technologies) and we verified all codon changes by DNA sequencing.

### Expression and purification of recombinant Hbs

We carried out recombinant Hb expression in the *E. coli* JM109 (DE3) strain as described previously (Natarajan et al., 2011). To ensure the complete cleavage of N-terminal methionines from the nascent globin chains, we over-expressed methionine aminopeptidase (MAP) by co-transforming a plasmid (pCO-MAP) along with a kanamycin resistance gene (Shen et al., 1993). We then co-transformed the pGM and pCO-MAP plasmids and subjected them to dual selection in an LB agar plate containing ampicillin and kanamycin. We carried out the over expression of each rHb mutant in 1.5 L of TB medium.

We grew bacterial cells at 37°C in an orbital shaker at 200 rpm until absorbance values reached 0.6 to 0.8 at 600 nm. We then induced the bacterial cultures with 0.2 mM IPTG and supplemented them with hemin (50 μg/ml) and glucose (20 g/L). The bacterial culture conditions and the protocol for preparing cell lysates were described previously (Natarajan et al., 2011). We resuspended bacterial cells in lysis buffer (50 mM Tris, 1 mM EDTA, 0.5 mM DTT, pH 7.0) with lysozyme (1 mg/g wet cells) and incubated them in an ice bath for 30 min. Following sonication of the cells, we added 0.5-1.0% polyethyleneimine solution, and we then centrifuged the crude lysate at 13,000 rpm for 45 min at 4°C.

We purified the rHbs by means of two-step ion-exchange chromatography. Using high-performance liquid chromatography (Äkta start, GE Healthcare), we passed the samples through a cation exchange-column (SP-Sepharose) followed by passage through an anion-exchange column (Q-Sepharose). We subjected the clarified supernatant to overnight dialysis in Hepes buffer (20 mM Hepes with 0.5mM EDTA, 1 mM DTT, 0.5mM IHP, pH 7.0) at 4°C. We used prepackaged SP-Sepharose columns (HiTrap SPHP, 5 mL, 17-516101; GE Healthcare) equilibrated with Hepes buffer (20 mM Hepes with 0.5mM EDTA, 1 mM DTT, 0.5mM IHP pH 7.0). After passing the samples through the column, we eluted the rHb solutions against a linear gradient of 0-1.0 M NaCl. After desalting the eluted samples, we performed an overnight dialysis against Tris buffer (20 mM Tris, 0.5mM EDTA, 1 mM DTT, pH 8.4) at 4°C. We then passed the dialyzed samples through a pre-equilibrated Q-Sepharose column (HiTrap QHP, 1 mL, 17-5158-01; GE Healthcare) with Tris buffer (20 mM Tris, 0.5mM EDTA, 1 mM DTT, pH 8.4). We eluted the rHb samples with a linear gradient of 0-1.0 M NaCl. We then concentrated the samples and desalted them by means of overnight dialysis against 10 mM Hepes buffer (pH 7.4). We then stored the purified samples at −80° C prior to the measurement of O_2_-equilibria and O_2_ dissociation kinetics. We analyzed the purified rHb samples by means of sodium dodecyl sulphate (SDS) polyacrylamide gel electrophoresis and isoelectric focusing. After preparing rHb samples as oxyHb, deoxyHb, and carbonmonoxy derivatives, we measured absorbance at 450-600 nm to confirm the expected absorbance maxima.

### Measurement of Hb-O_2_ equilibria

Using purified rHb solutions (0.3 mM heme), we measured O_2_-equilibrium curves at 37°C in 0.1 M Hepes buffer (pH 7.4) in the absence (‘stripped’) and presence of 0.1 M KCl and IHP (at two-fold molar excess over tetrameric Hb), and in the simultaneous presence of KCl and IHP. We measured O_2_-equilibria of 5 μl thin-film samples in a modified diffusion chamber where absorption at 436 nm was monitored during stepwise changes in the equilibration of N_2_/O_2_ gas mixtures generated by precision Wősthoff mixing pumps (Weber, 1992; Grispo et al., 2012; Weber, Fago, Malte, Storz, & Gorr, 2013). We estimated values of *P*_50_ and *n*_50_ (Hill’s cooperativity coefficient) by fitting the Hill equation *Y*= PO_2_^*n*^/(*P*_50_^*n*^+PO_2_^*n*^) to the experimental O_2_ saturation data by means of nonlinear regression (*Y* = fractional O_2_ saturation; n, cooperativity coefficient). The nonlinear fitting was based on 5-8 equilibration steps. Free Cl^−^ concentrations were measured with a model 926S Mark II chloride analyzer (Sherwood Scientific Ltd, Cambridge, UK).

### Measurement of Hb-O_2_ dissociation kinetics

We determined O_2_ dissociation constants (*k*_off_) of purified rHbs at 37 °C using an OLIS RSM 1000 UV/Vis rapid-scanning stopped flow spectrophotometer (OLIS, Bogart, CA) equipped with an OLIS data collection software. Briefly, rHb (10 μM heme) in 200 mM Hepes, pH 7.4, was mixed 1:1 with N_2_-equilibrated 200 mM Hepes, pH 7.4, containing 40 mM sodium dithionite (Helbo& Fago, 2012). We monitored absorbance at 431 nm as a function of time and fit the curve to a monoexponential function (*r*^2^ > 0.99). We calculated O_2_ association rates (*k*_on_) from the relationship *k*_on_ = *k*_off_/*K*, where *K* (μM) is the O_2_ dissociation equilibrium constant in solution, calculated as the product of *P*_50_ (torr) and the O_2_ solubility coefficient in water at 37° C (1.407 mM torr^−1^)(Boutilier, Heming, & Iwama, 1984).

### Measurement of autoxidation rates

To estimate autoxidation rates, we treated purified rHb samples with potassium ferricyanide (K_3_[Fe(CN)_6_]), and we then reduced rHbs to the ferrous (Fe^2+^) state by treating the samples with sodium dithionite (Na_2_S_2_O_4_). We removed the dithionite by means of chromatography (Sephadex G-50). For each rate measurement, we used 200 μl of 20 μM oxyHb in 100 mM potassium phosphate buffer, pH 7.0, containing 1 mM EDTA and 3 mM catalase and superoxide dismutase per mole oxyHb. To measure the spontaneous conversion of ferrous (Fe^2+^) oxyHb to ferric (Fe^3+^) metHb we recorded the absorbance spectrum at regular intervals over a 90 h period. We collected spectra between 400nm and 700nm using a BioTek Synergy2 multi-mode microplate reader (BioTek Instruments). We estimated autoxidation rates by plotting the A_541_/A_630_ ratio (ratio of absorbances at 540nm and 630nm) vs time, using IGOR Pro 6.37 software (Wavemetrics). We used the exponential offset formula in IGOR to calculate the 50% absorbance per half-life (i.e., 0.5AU/half-life).

### Measurements of structural stability

We assessed the pH-dependent stability of the rHbs by means of UV-visible spectroscopy. We prepared 20 mM filtered buffers spanning the pH range 2.0-11.0. We prepared 20 mM glycine-HCl for pH 2.0–3.5; 20 mM acetate for pH 4.0–5.5; 20 mM phosphate for pH 6.0–8.0; 20 mM glycine-NaOH for pH 8.5–10.0; 20 mM carbonate-NaOH for pH 10.5 and phosphate-NaOH for pH 11.0. We diluted the purified rHb samples in the pH-specific buffers to achieve uniform protein concentrations of 0.15 mg/ml. We incubated the samples for 3–4 h at 25°C prior to spectroscopic measurements, and we maintained this same temperature during the course of the experiments. We measured absorbance in the range 260–700 nm using a Cary Varian Bio100 UV-Vis spectrophotometer (Varian) with Quartz cuvettes, and we used IGOR Pro 6.37 (WaveMetrics) to process the raw spectra. For the same set of rHbs, we tested for changes in secondary structure of the globin chains by measuring circular dichroism spectra on a JASCO J-815 spectropolarimeter using a quartz cell with a path length of 1 mm. We assessed changes in secondary structure by measuring molar ellipticity in the far UV region between 190 and 260 nm in three consecutive spectral scans per sample.

### Structural modeling

We modelled structures of goose Hbs and the various mutational intermediates using the program COOT (Emsley, Lohkamp, Scott, & Cowtan, 2010), based on the crystal structures of bar-headed goose Hb (PDB models 1hv4 and 1c40)(Liang, Hua, et al., 2001; Liu et al., 2001), greylag goose Hb (PDB 1faw)(Liang, Liu, Liu, & Lu, 2001), and human deoxyHb (PDB 2dn2).

## Acknowledgements

We thank E. E. Petersen for skilled assistance, H. Moriyama for sharing equipment, and K. G. McCracken for helpful discussion. This work was funded by the National Institutes of Health/National Heart, Lung, and Blood Institute (HL087216 [JFS]), the National Science Foundation (MCB-1517636 [JFS] and RII Track-2 FEC-1736249), and the Danish Council for Independent Research, Natural Sciences (4181-00094 [AF]).

---

**Figure supplement 1**. Effects of forward and reverse mutations on the same backgrounds.

**Figure supplement 2**. Variation among goose rHb mutants in functional and structural properties that potentially trade-off with intrinsic O_2_ affinity.

**Source data 1**. Autoxidation rates of all rHb mutants.

**Source data 2**. Effect of pH on the stability of α-helical secondary structure of rHb mutants, measured by circular dichroism spectroscopy.

**Source data 3**. Effect of pH on the stability of tertiary structure of rHb mutants, measured by UV-visible spectroscopy.

**Figure supplement 1.** Ramachandran plot of deoxyHb from bar-headed goose (PDB 1hv4).

**Figure 3 - figure supplement 1.**
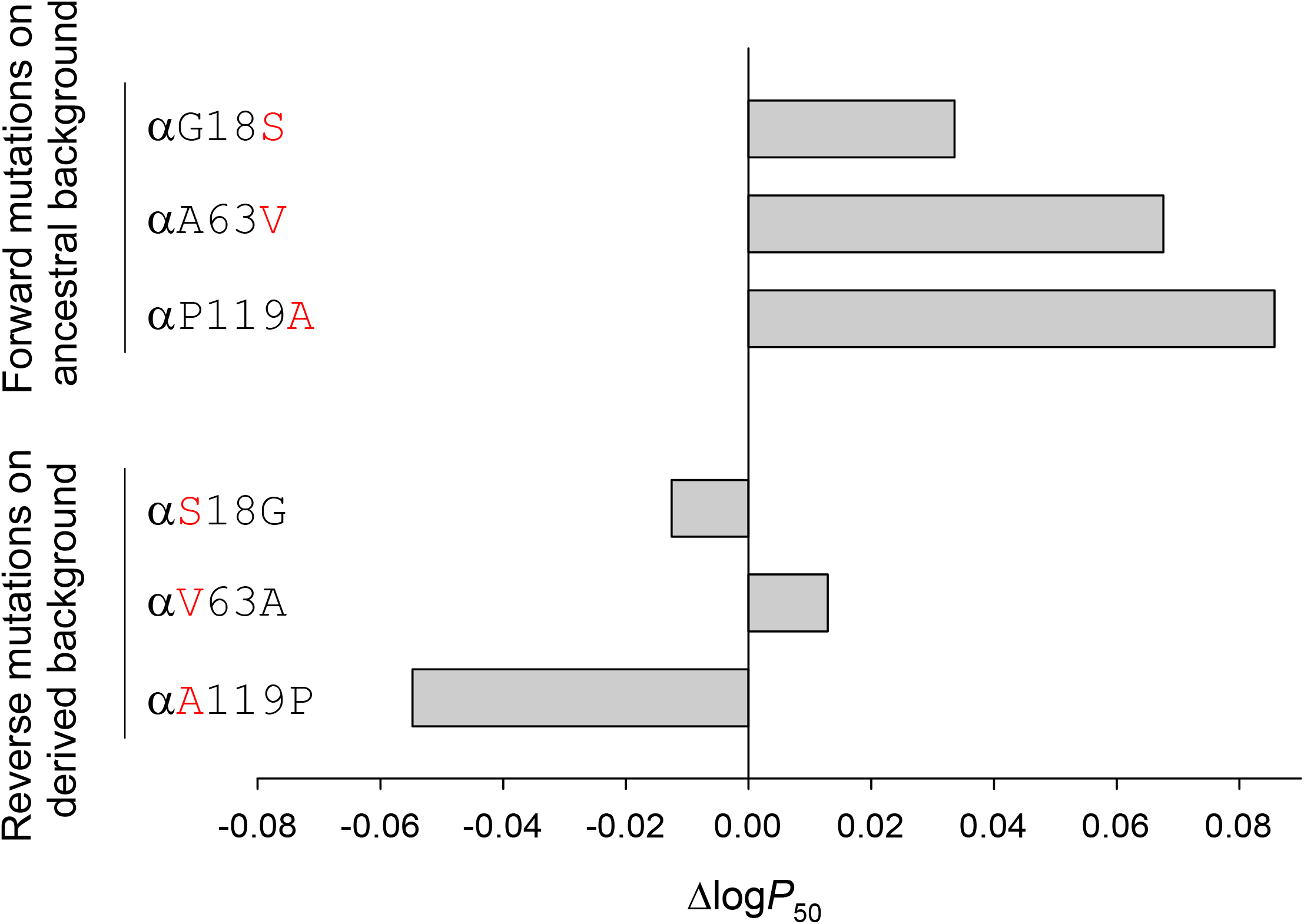
The αP119A mutation has consistent effects on Hb-O_2_ affinity on different genetic backgrounds. The affinity-enhancing effect of αP119A on the AncAnser background is mirrored by a similarly pronounced affinity-reducing effect when the mutation is reverted on the wildtype bar-headed goose background (αA119P). By contrast, forward and reverse mutations at α18 and α63 do not show the same symmetry of effect, indicating that their effects are conditional on the amino acid state at one or both of the other two sites.

**Figure 3 - figure supplement 2.**
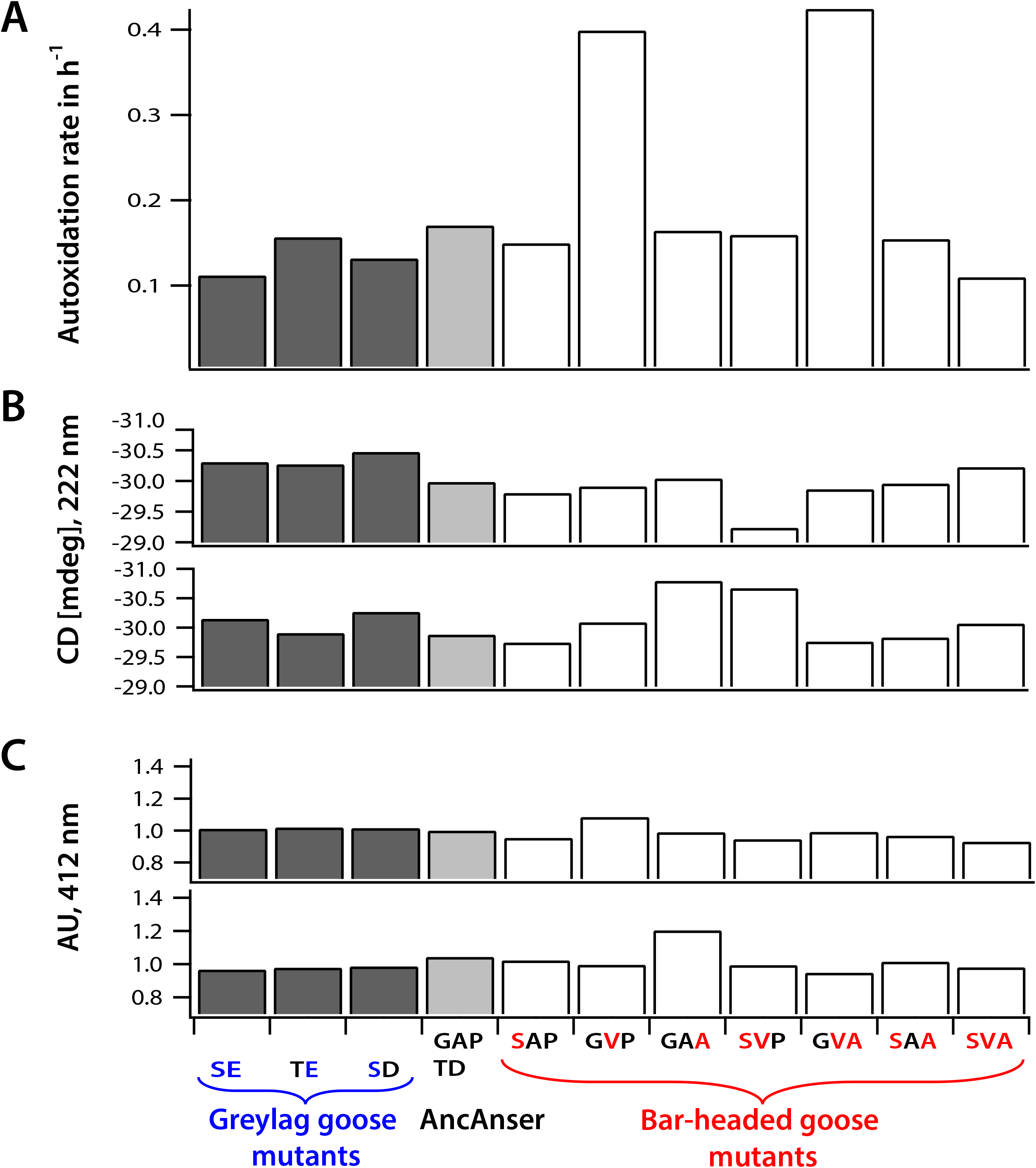
Variation among goose rHb mutants in functional and structural properties that potentially trade-off with intrinsic O_2_ affinity. Variation in (*A*) autoxidation rate (rate at which ferrous heme [Fe^2+^] spontaneously oxidizes to the ferric state [Fe^3+^]), (B) stability of secondary structure, as assessed by means of circular dichroism spectra (with ellipticity measured in millidegrees [mdeg], 222 nm) at pH 7.0 and 7.5 (physiological range), and (C) stability of tertiary structure and holoprotein, as assessed by means of UV-visible spectroscopy (absorbance measured at 412 nm) at pH 7.0 and 7.5 (physiological range). For stability measurements over the full pH range, see Data Source Files 1-2.

**Figure 4 - figure supplement 1.**
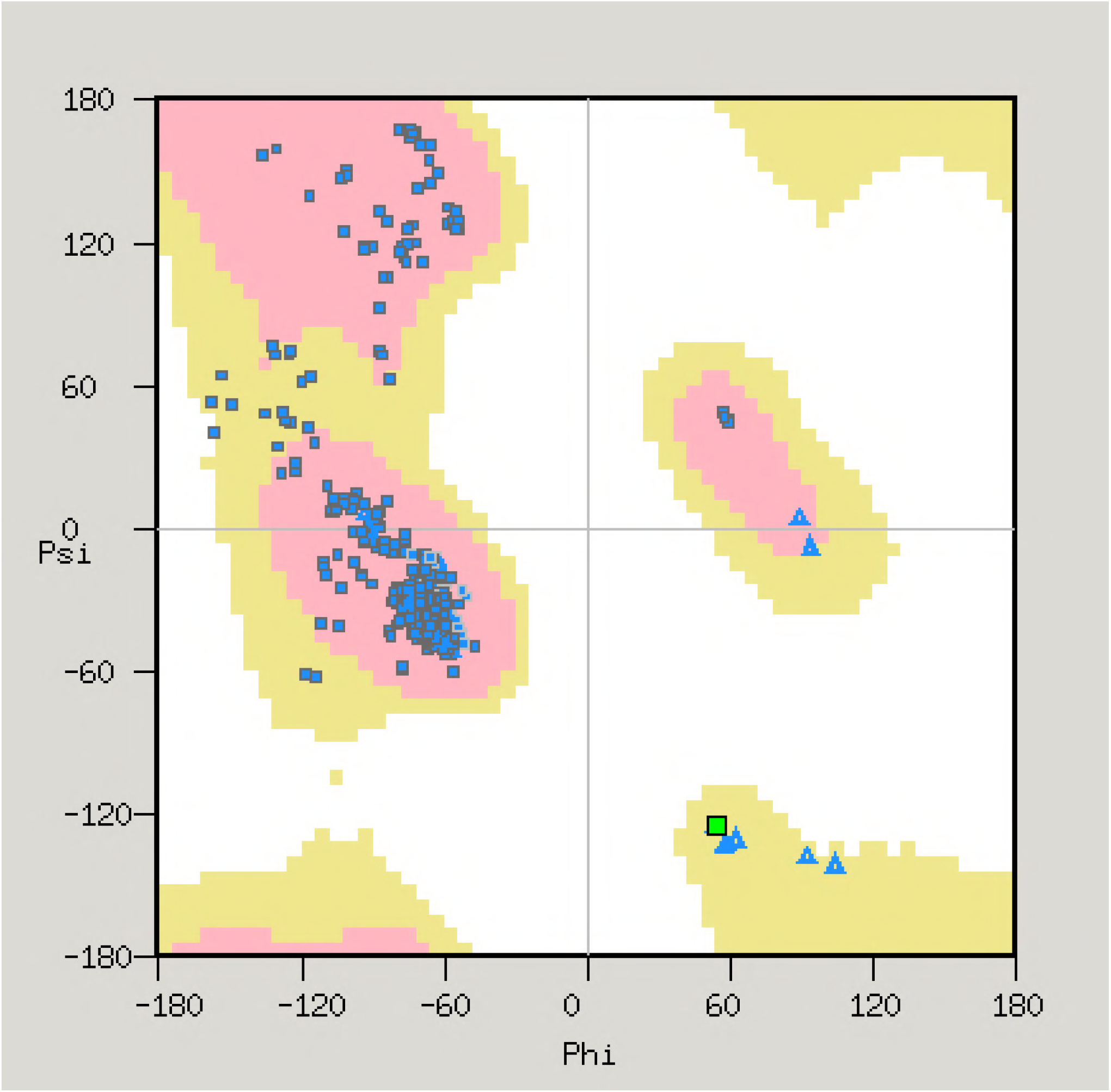
Ramachandran plot of deoxyHb from bar-headed goose (PDB 1hv4). Glycine residues are denoted by triangles, other residues by squares. One residue, α18-Ser, is conspicuous by its unusual backbone angles, and is shown as a green square. This position in the Ramachandran plot is highly unusual for any residue other than glycine. The turn in the backbone between the A and B helices can only be accommodated by a glycine, since the lack of a side-chain avoids the strong steric clash that would develop between a C_β_ atom and the nitrogen atom of residue 19. The serine at α18 is therefore forced to flip the peptide conformation, such that its carbonyl group points in the opposite direction relative to that of Gly 18.

